# Introducing CARATE: Finally speaking chemistry through learning hidden wave-function representations on graph-based attention and convolutional neural networks

**DOI:** 10.1101/2022.02.12.470636

**Authors:** Julian M. Kleber

## Abstract

Computer-aided drug design is stepping into a new era. Recent developments in statistical modelling, including *deep learning, machine learning* and high-throughput simulations, enable workflows and deductions unachievable 20 years ago. The key interaction for many small molecules in the context of medicinal chemistry is via biomolecules. The interaction between a small molecule and a biological system therefore manifests itself at multiple time and length scales. While the human chemist may often grasp the concept of multiple scales intuitively, most computer technologies do not relate multiple scales so easily. Numerous methods that try to tackle multiple scales in the realm of computational sciences have been developed. However, up to now it was not clear that the problem of multiple scales is not only a mere issue of computational abilities but even more a matter of accurate representation. Current representations of chemicals lack the descriptiveness necessary for today’s modelling questions. This work introduces a novel representation of small and large molecules. The representation is obtained by the novel *biochemical and pharmaceutical encoder* (CARATE). In the following work, the regression and classification abilities of the learned representation by CARATE are evaluated against benchmarking datasets (ZINC, ALCHEMY, MCF-7, MOLT-4, YEAST, ENZYMES, PROTEINS) and compared to other baseline approaches. CARATE outperforms other graph-based algorithms on classification tasks relating to large biomolecules and small molecules, as well as on quantum chemical regression tasks of small molecules.

## 1 Introduction

The wish to accurately predict the outcome of a drug development process is as old as the regulated pharmaceutical industry itself. However, the task of simulating a drug interaction in the body is a multiscale problem operating on more than two scales.

Up to now, no accurate method has been found to model the whole life cycle of an active pharmaceutical ingredient (API) inside a mammal. The general work on multiscale problems focuses on different aspects of the systems at multiple scales and strives to represent the relevant aspects of a specific scale accurately by means of that particular scale. The simulator then decides on apparently unimportant parts of that scale and approximates them more efficiently by employing methods more commonly used at a larger scale ^1^.

The problem of multiple scales shows us that our current level of theory does not handle the notion of multiple scales well enough to match our computational abilities. Therefore, each multiscale modelling attempt is doomed to fail using our current technology alongside our current state of theoretical physics describing the dynamics of the system.

Because of the number of already performed simulations at various scales, deep learning (DL) becomes a viable approach for speeding up computations by approximating simulations in a data-driven way.

Yet the current representation of chemicals (SMILES, InChI and Lewis-like structures) cannot be suitable for modelling because they are *themselves* abstractions. Even more problematically, the original SMILES algorithm was claimed to be unique ^2^, but was later proven to be non-unique ^3^. It is impossible for fingerprints and SMILES strings to represent a wave function properly and so methods using these representations are doomed to fail.

The recent surge in application of SMILES for use in DL ^4–9^ leaves doubts about the applicability of the derived models. On the contrary, large matrix representations are mostly inaccurate while being resource intense ^10^. To accurately approximate the physical (simulated) properties of a given compound, a more accurate, uniform representation is necessary.

When different length and time scales are involved, computationally expensive methods for the small-length scale may become infeasible. For example, to model the interaction of a small molecule with a receptor, at the moment one needs to combine quantum chemistry (QC) with classical molecular mechanics (MM).

The computations performed by the QM/MM approximation still have the QC calculations as a bottleneck. However, recent advances in DL are making the prediction of a particular simulation scale accessible ^11,12^. Recently the prediction of quantum chemical properties with deep-learning methods became more accurate ^13^ leading to MD simulations using learned coupled-cluster potentials ^14^.

Implicit methods might thus solve the equation much more efficiently and accurately. To speed up the computation of a QM/MM simulation, the QC computations should be approximated by predicting the desired properties via machine learning methods ^15^.

There are several exceptional graph neural networks reported on the PROTEINS and ENZYMES dataset which have gained good scores ^16–18^. The most prominent ones are hierarchical structure methods employing convolutional layers as well as message-passing layers ^19,20^.

The clustering methods have also been achieving high accuracies on the PROTEINS and ENZYMES data set (no test set), and high accuracies on the QM family have been reported ^20^.

For example, *Chen et al*. report a similar algorithm to CARATE (this work) developed during the same time, but omit results on large datasets, which they used to pretrain their algorithm ^9^.

Moreover, there is a report of a hyperparameter scan for meta-learning graph models that shows a good fitting of the PROTEINS dataset of 73.8% ^21^. However, the performance does not seem to be satisfactory.

Moreover, in the original GAT paper, the authors did not fit the algorithm on either of the ENZYMES or PROTEINS datasets although they achieved good results on the protein-interaction dataset (PPI) ^22^. The PPI data set, however, is minimal and may not be representative for any fitting behavior.

Yet graph-based methods do not reach the accuracy necessary for accurate simulation because all methods neglect self-interaction, a foundational quantum mechanical principle. Either the graph-based algorithm ignores self-interaction, or the given input neglects self-interaction.

The current graph-based methods thus lead to an already flawed ground truth and are doomed to fail. Therefore, what most methods have in common is that they try to incorporate quantum-chemical principles but are missing the self interaction terms.

Presently, it is barely possible to perform simulations of decent size and complexity on edge devices or personal computers. Typically, the equipment involved consists of nodes on a high-performance cluster. The fact that only insiders can access high-performance clusters gives rise to an ever more arcane elite of practitioners. A problem chemistry was facing since its birth in alchemy.

A novel method equipped with algorithms of outstanding performance might lead to a new era of edge simulations to enable scientists, as well as to accelerate and democratize progress in the natural and life sciences.

Until now, however, to the best of my knowledge, no algorithm with short training times and competitive prediction abilities on edge devices has been reported.

This work investigates whether deep-learning methods can be exploited to solve the time-independent Schrödinger equation instead of bypassing it via predictions. The working idea is to use molecular graphs, encode them into a wave function and decode them subsequently such that the eigenvalue equation 2 is solved.

Under the hood, molecular graphs are adjacency matrices that can be equipped with more features through multiple annotation matrices, such that the graph representation carries much more information than plain SMILES strings.

The problem then looks similar to a classical deep-learning method, framing the families of the eigenvalue problems described in equation 4 as a problem that can be tackled via deep learning.

This work aims to close the gap by providing a robust method that makes sure, that the representation, the neural-network, and the training process obey quantum-physical principles.

## 2 Related Work

Predicting physical properties of chemical compounds with statistical methods is indeed a well-known topic in the literature. Recent work comparing the representations of molecules for the QM-9 prediction performance of a feed-forward neural network (FNN) showed that the quality of predictions of physical properties depends strongly upon the molecular representation ^23^.

The recent advances in natural language processing (NLP) introduced by the self-attention mechanism in bidirectional auto encoders ^24^ showed that respective algorithms (BERT technology) can learn semantic properties of the input ^25^. In particular, the BERT algorithm has abilities to organize and structure information ^26^.

Molecular descriptors can be learned in a data-driven way ^27^. Therefore, the self-attention mechanism made room for innovation around self-attention deep-neural networks in computational chemistry using the SMILES or similar text representations of molecules with good classification and regression results ^28^.

Indeed, semi-empirical potentials are well-known to the community of computational chemists. Especially, the extended tight-binding methods are fitted in a statistical manner from a large set of molecules and have been celebrating a huge success ^29^.

Thus, to the community, it does not seem to be relevant why something works as long as it works. In contrast to most semi-emprical potentials, deep-learning methods offer explainability through dedicated algorithms such as LRP ^30^ and others.

A fundamentally different, yet equally promising approach to a text-based encoding is to use numerical representations such as fingerprints, substructures, or graphs. Indeed, there is a recent study investigating the potential of unified fingerprints for small molecules as well as biomolecules ^31^.

However, one-shot learning based on self-attention LSTM with encoded molecular graphs as inputs showed more decent results ^32^.

Many approaches for modelling the molecular structure with graph neural networks (GNNs) have been applied. To increase computational efficiency, a self-attention mechanism has been rudimentarily applied to graph convolutional networks (GCNs( as well ^28^.

In 2017, the group around *Schütt et al*. proposed *SchNet* and their benchmark dataset *ISO17* ^33^. The key improvement in *SchNet* was to capture the molecular geometry as a graph while using convolutions to craft weighted feature importance in the molecule.

Another way to achieve better structure recognition is to apply message-passing networks ^34^. Furthermore, multi-headed self-attention has been applied to GNN-encoded molecular structures through a self attention mechanism ^35^.

*Chen et al*. introduced the idea of bidirectional transformers to chemistry on graphs. Yet they still work with SMILES and also have not published their results on large benchmark datasets ^9^. The idea of obtaining a better representation through graph-based deep-learning is becoming more and more popular; however, many chemists are still working on SMILES representations ^20^.

This work completes the series on the application of attention-based algorithms to chemical problems. It investigates a multi-headed graph self-attention model (GAT) ^22^ with graph-convolutions as an encoder for chemical classification and regression tasks, with significant results.

## 3 Theoretical Remarks

For quantum chemistry at any scale, we want to solve the time-dependent Schrödinger equation when considered in the context of the dynamic behaviour of the system.

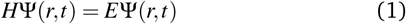

As elaborated above, one of the working hypotheses is that the current machine-learning methods, including modern deep-learning methods, in general only bypass the time-independent Schrödinger equation by making predictions.

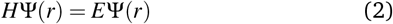

But what does it mean to solve equation 2? We are generally interested in finding eigenvalues and eigenfunctions of a system for a particular operator (in this case, the Hamiltonian). Very often, only the eigenvalues are of interest.

The solution to the time-dependent part of equation 1 is, however, rather simple. First one starts by separating the variables

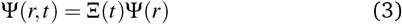

We then solve for the positional part and the time-dependent part separately. The time-dependent part has standard solutions that have an analytical expression.

The time-independent part, however, has solutions that are not solvable by analytical means for many-electron systems. The only pathway to a solution for the time-independent part of the Schrödinger equation is thus a numerical one.

The accuracy of each method for solving equation 2 is thus theoretically bounded by the hardware limits of the machine performing the calculations.

The more interesting problem is to solve the *time-dependent* Schrödinger equation. Once the TISE is solved, the dynamics can be computed efficiently.

Solving the TISE with deep learning is not a new idea and has been prominent on tech blogs since 2017^36^. The article by *Steinke* provides a first hint that the problem is tractable with neural networks.

In the academic literature, the topic occurs in relatively frequently just a little time later ^37–40^. However, the methods presented by the authors differ from this paper fundamentally by explicitly incorporating the flaws and approximations that classical quantum chemistry must make by design.

The key innovation in algorithms exploiting deep learning is not backpropagation, but automatic differentiation (AD). It has been shown that AD can be exploited to tackle quantum chemical calculations because with AD it is possible to optimize basis sets in the Hartree-Fock scheme ^41^. Moreover, the application of physical regularizers has been shown to be beneficial in modelling wavefunctions via backpropagation ^42^.

The approximations and flaws include the Born-Oppenheimer-approximation (BOA), using Kohn-Scham density functionals instead of wave functions, and the approximation of the wave function as an expansion of an infinite basis set.

The standard expansion of the wave function into atomistic basis functions needs numerous determinant calculations (known as Slater determinants) and are thus lengthy to perform. The self-critical comments on FermiNet conclude that the scalability of FermiNet might not be useful for practical quantum chemistry ^37^.

Moreover, many authors ignore the fact that most quantum chemical methods, including the coupled-cluster methods, rely on the Hartree-Fock optimized basis sets as their starting point.

In the end, all quantum chemical methods want to solve the eigenvalue equation 2. It is thus mathematically always the same problem, and just the direction from which it is tackled differs.

The creativity in problem-solving comes thus from solving the general equation for arbitrary observables

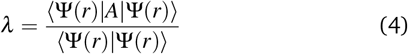

In most cases, chemists are not interested in the wave function nor how it has been constructed, and an explicit calculation might not always be necessary.

Any explicit calculation focusing on explainability of the employed basis sets might even lead to misleading concepts that chemists then apply in other disciplines without having any means of thinking critically about the applied models.

For example, the orbital concept is a simplification that comes from the aforementioned approximations made by theoretical chemists and was made popular during the 20th century in the western community, especially, by Linus Pauling.

Most ironically, Linus Pauling also introduced the construction of arbitrary wave functions through combining atomistic wave functions into hybrid orbitals (molecular orbitals) using group theory in his valence bond theory.

Today, most (quantum) chemistry still builds on this idea. The general quantum-chemical calculation is built up through calculating better wave functions obtained from simpler wave functions. Equation 5 demonstrates the principle of building wave functions of higher scale from wave functions of lower scales ^43^. The order of building the wave function starts from atomic orbitals (AO), and goes over molecular orbitals (MOs), Slater determinants (SD) and finally many-electron wave functions (ME).

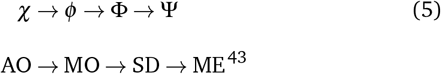

Starting from the idea of constructing hybrid wave functions, this approach is not far off taking other features and combining them into even more sophisticated wave functions that are then combined into a system-wide wave function.

The concept of constructing a holistic wave function directly from diverse input features or smaller wave functions is inherently explored in this work by applying the CARATE algorithm to the datasets ALCHEMY and ZINC. Both of these datasets have a diverse set of features.

Thus, this works aims to construct Ψ directly from the annotated molecular graph, taking the annotated graph nodes and edges as building blocks for the wave function Ψ.

Thereby, the new method CARATE shall skip at least two steps in equation 5 and be independent of the BOA. Further, the new CARATE method does not rely on Euclidean coordinates and thus operates in a space other than the Euclidean space. The method does not rely on Hartree-Fock calculations but uses the variational principle by default.

## 4 Methods

The whole code was forged into a framework called CARATE and moved later to a package called *aiarc*. Both packages are open source and available on Codeberg, a German association for the advancement of open-source software (https://codeberg.org/sail.black/carate). The package can be obtained via PyPi and pip, too (https://pypi.org/project/carate/).

The package contains a documentation hosted on readthedocs.io as well as notebooks providing the analysis of the training trajectories and some tutorials on how to start a run from a notebook.

The configuration files are provided as supporting information, as well as extra plots, trajectories, log files, and trained models. The reader is referred to the Codeberg of the package to obtain the supporting information.

Moreover, the QM9 dataset was omitted because the proof of concept which entailed solving the problem presented by QM9 was already tackled via the ALCHEMY dataset. The ALCHEMY dataset even presents a more difficult task, as there are more heavy atoms per molecule on average than in the QM9 dataset.

In general, the training on GPU is slightly faster, though the CPU was chosen as the reference chip. The accuracy of the model is sometimes slightly lower and less reproducible across different CUDA versions and GPUs at different times.

For the presented work, an AMD Ryzen 9 7950X3D at 4.5 GHZ 32 Core, and AMD Ryzen 9 7950X processor 16Core at 4.5 GHz and 64 MB L3 Cache alongside 64 GB of RAM were used on a Debian 12 Bookworm system.

Different systems were tested during the development and all produced the same results. Among the tested systems were Ubuntu 20.04, 22.04, Debian 11, EndeavorOS, and Debian 12. The algorithms were reproducible on all of them. Thus, both the Arch Linux and common Debian distributions are suitable for modelling tasks associated with the package.

Both methods, regression and classification, use the basic encoder structure and apply specific architectures on top of the encoder structure for each particular task. For classification, the outputs of the encoder were transformed into logits using a sigmoid function. For regression, a linear layer was stacked on top of the encoder structure.

To test the classification abilities, the model was tested against the drug discovery datasets for small molecules YEAST, MOLT-4, and MCF-7 obtained from the TUDataset without any further preparation (Table 4).

To test the performance on large protein graphs, the ENZYMES as well as the PROTEINS datasets were chosen. The datasets were obtained from the TUDataset using the PytorchGeometric API without any further preparation.

For each example, five-fold stratified cross-validation for statistical significance of the results was performed.

The datasets were obtained from TUDataset ^44^, and MoleculeNet ^45^ through the PyTorchGeometric API ^46^. Each dataset was used as is, to see how CARATE performs on raw data. Only the regression datasets were normalized to compare them to the baseline approaches. Thus, the only preprocessing applied was rescaling to the target’s unit for regression tasks, as would be done during a simulation.

If the data set were given in SMILES format, the SMILES format was then converted to a molecular graph representation implemented in PyTorchGeometric ^46^. All GAT and GNN implementations were applied from or achieved with the PyTorchGeometric ^46^ library. Other deep-learning building blocks were implemented using the PyTorch ^47^ and Numpy ^48^ libraries.

The CARATE network consists of an input layer consisting of a GCN and GAT layer. To use the encoder in regression and classification tasks, another GCN layer followed by task-specific layers are added to the stack (compare Figure 1)

**Fig. 1.**
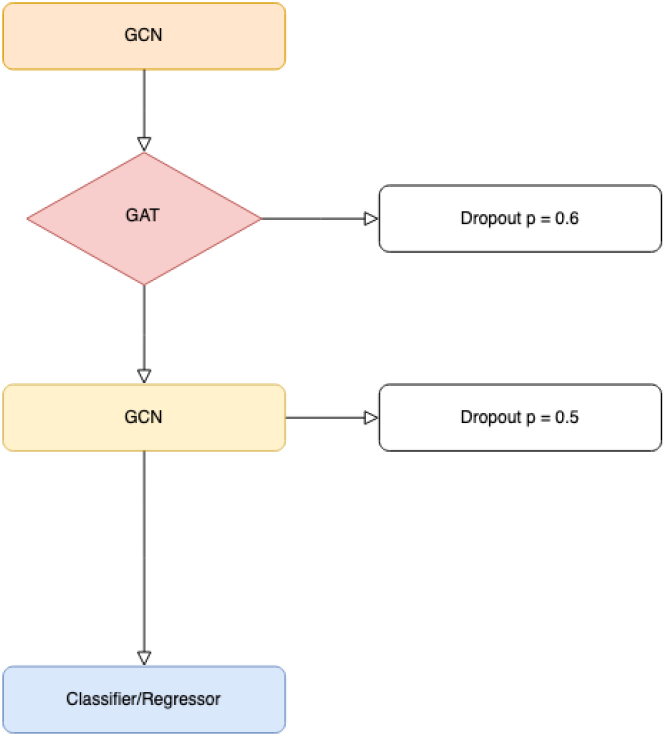
General architecture of the CARATE network used for classification and regression tasks.

To test the encoding to property prediction, the encoder-predictor model was tested against physical regression tasks on the ALCHEMY ^49^ and ZINC datasets obtained from TUDatasets ^44^.

The architecture for the regression tasks differs from the classification to obtain multitask regression results, instead of multitask classification only by the output function applied to the network.

The results are compared to GIN-*ε* ^44,50^.The GINE-*ε* network uses batch normalization as well as zero mean and unit variance normalization.

### Ablation studies

To examine the importance of each building block, an ablation study was performed. As reference datasets, the ALCHEMY and ZINC datasets for regression were used, as well as the PROTEINS and ENZYMES dataset for classification.

The tested simplified algorithms included: just a GAT unit, just a convolutional unit, the encoder GCN-GAT-GCN without the two linear layer operators, and finally just the linear units used as the operator to act on Ψ.

To enable fitting with just linear networks work, a pooling layer was introduced to account for the mini batch. This is rationalized as, in general, GNNs are a generalization of GCNs and adding a pooling layer would be necessary for a classical GCN as well.

Proper convergence steps were selected on the previous experiments for the CARATE algorithm. The regression runs were performed five times and ran for ten training steps. The classification tasks were performed five times, and ran for 2000 steps each.

## 5 Results

### CARATE

The reader with historical interest can find the original notebooks containing the first experiments are in the Codeberg: (https://codeberg.org/cap_jmk/carate-notebooks)

The convergence is for each task quite fast and well below 1000 iterations.

In fact, early stopping was beneficial in most cases, such that longer training results may lead to less accurate results, especially for the classification tasks. The described degradation of the convergence behaviour indicates that the algorithm is prone to over-fitting.

For the small-molecule regression tasks consisting of the ZINC and the ALCHEMY datasets, the training converged after less than 10 epochs (usually two epochs) up to numerical accuracies (Table 2. There is a slight dependence on the train-test split (Figures 2,3). The protein classification tasks consisting of the ENZYMES and PROTEINS datasets showed exceptional results that outperformed all reported benchmarks (Table 4).

**Table 1.** Results of the ablation study.

| Method | ZINC 500k (Accuracy) | Alchemy 500k (Accuracy) | Enzymes 1.2M (MAE) | Proteins 1.2M (MAE) |
| --- | --- | --- | --- | --- |
| GATv2 | $0.003 \pm 0.002$ | $0.001 \pm 0.001$ | $26 \pm 5$ | $78 \pm 6$ |
| GCN | $0.03 \pm 0.05$ | $0.1 \pm 0.2$ | $13 \pm 5$ | $61 \pm 3$ |
| GCN + GAT | $0.02 \pm 0.02$ | $0.03 \pm 0.05$ | $8 \cdot 10^1 \pm 1 \cdot 10^1$ | $89 \pm 3$ |
| Linear + Pooling | $0.1 \pm 0.2$ | $0.06 \pm 0.1$ | $16 \pm 4$ | $44 \pm 3$ |
| CARATE one linear | $0.003 \pm 0.002$ | $0.001 \pm 0.002$ | $9 \cdot 10^1 \pm 1 \cdot 10^1$ | $96 \pm 6$ |

**Table 2.** Regression performance compared to baseline GINE-*ε* ^44^.

| Method | ZINC 500k | ALCHEMY 500k |
| --- | --- | --- |
| CARATE | $0.0016 \pm 0.0002$ | $0.00050 \pm 0.00005$ |
| GINE-ε<sup>44</sup> | 0.084 | 0.103 |

**Fig. 2.**
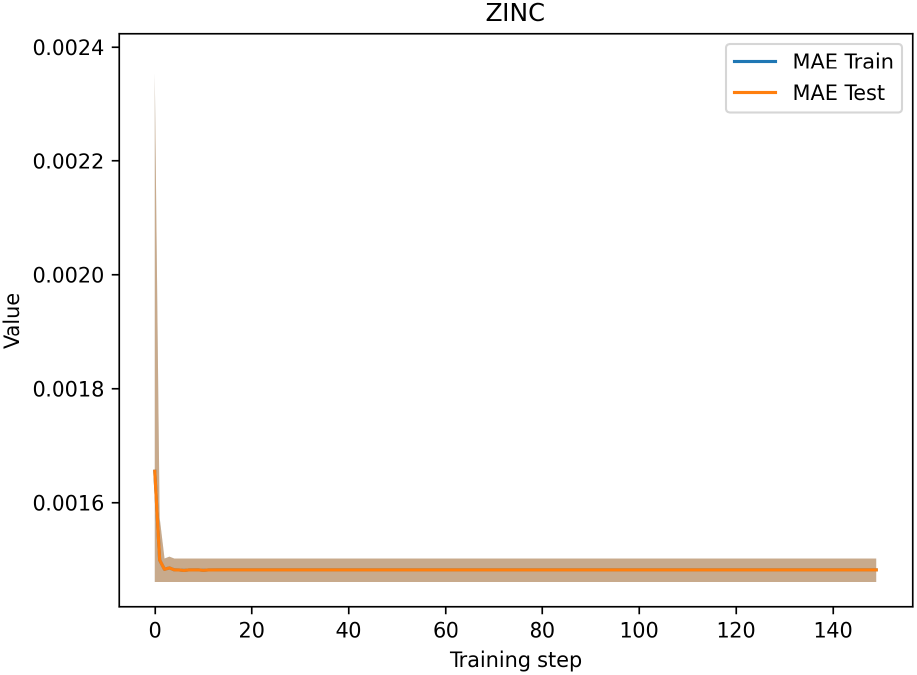
Accuracy trajectories for both training and test sets for *n* = 5000 training steps on the ZINC dataset.

**Fig. 3.**
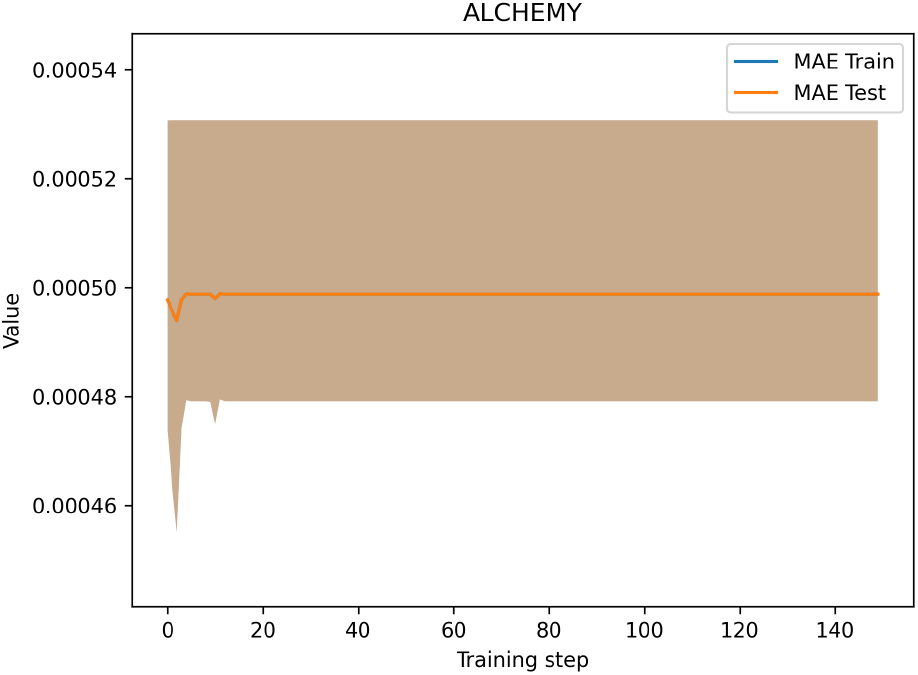
Accuracy trajectories for both training and test sets for *n* = 5000 training steps on the ZINC dataset.

On both protein classification tasks, ENZYMES and PROTEINS, CARATE provides best-of-class fits, with accuracies up to 100% on certain splits (compare Figures4, 5). The algorithm was prone to slight overfitting on both datasets.

**Fig. 4.**
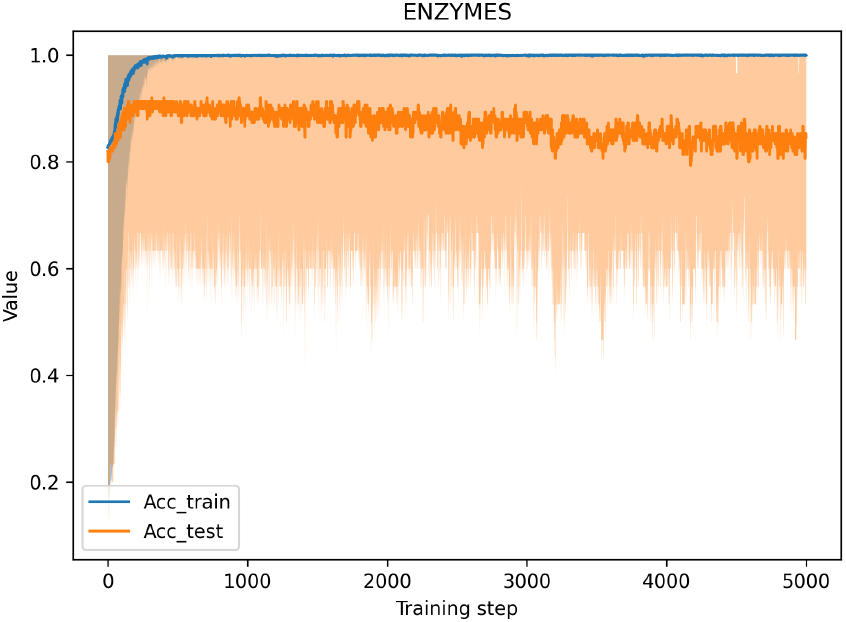
Accuracy trajectories for both training and test sets for *n* = 5000 training steps on the ENZYMES dataset.

**Fig. 5.**
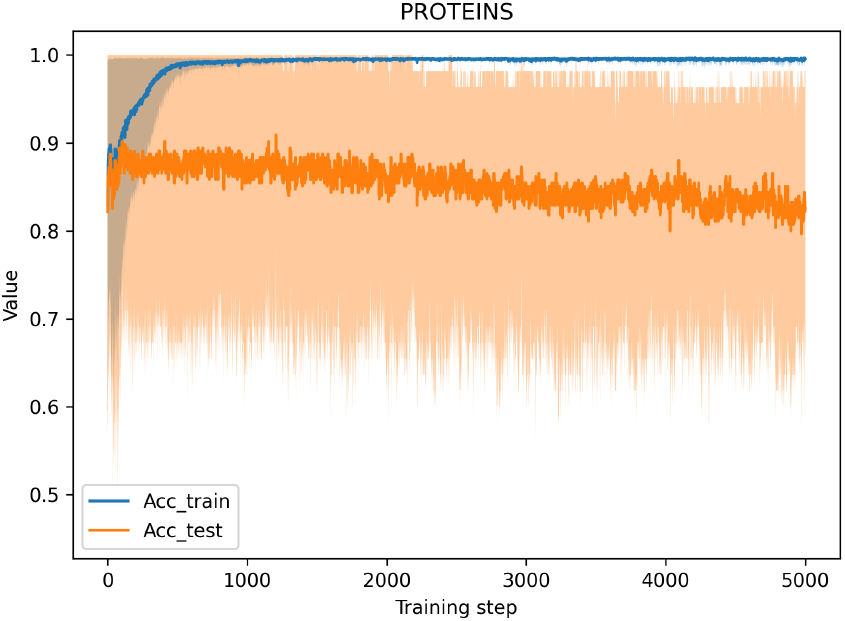
Accuracy trajectories for both training and test sets for *n* = 5000 training steps on the PROTEINS dataset.

**Fig. 6.**
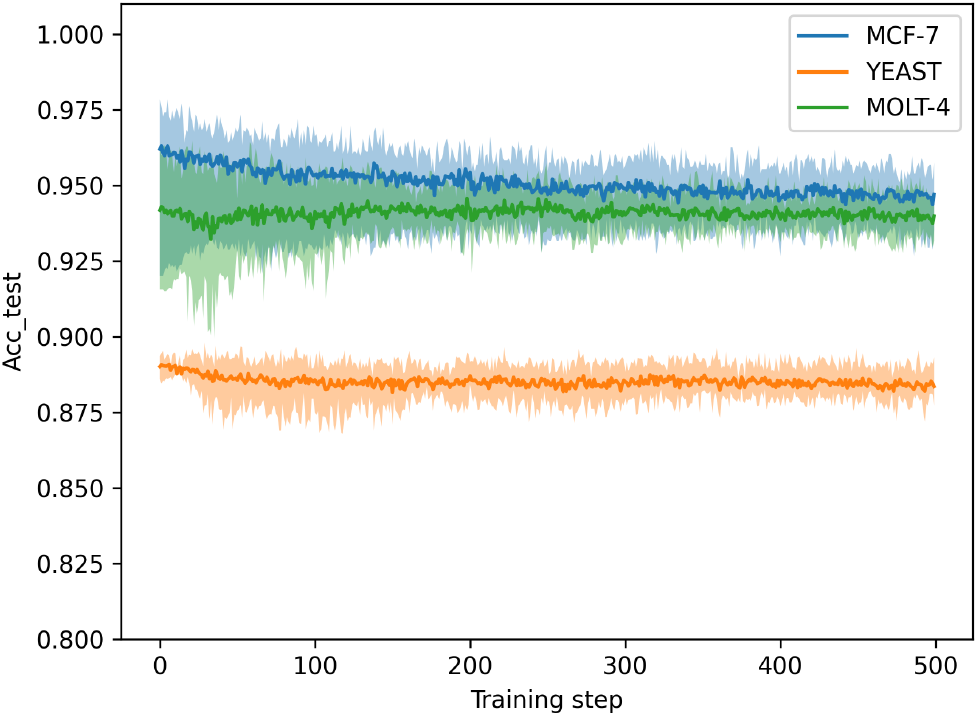
Trajectory of the training accuracy using the datasets MCF-7, MOLT-4, and YEAST.

### Ablation Study

The results of the ablation study are shown in 1. The ablation study strongly suggests that neither part of the CARATE algorithm can achieve full performance on its own. Yet, for the CARATE algorithm, the second linear layer does not improve the accuracy of the algorithm much, but only reduces variance.

## 6 Discussion

The aim of efficiently learning a decent molecular to property representation from a dataset has been achieved. The proposed architecture can be applied to small and large biomolecules alike with satisfying results. The CARATE encoder enables multi-class classification and regression tasks.

In Table 4, CARATE is compared to several other state-of-the-art methods in the field. SchNet ^33^ is based on the findings of DTNN ^51^. Thus, DTNN is one of the foundational works and has hardly ever been matched by subsequent works in explicit wave function modelling. With DTNN *Schütt et al*. investigated the use of deep learning to solve the many-body Schrödinger equation. Subsequent work such as PauliNet ^38^ and SchNorb ^52^ further build upon the idea.

**Table 3.** **Comparison of different algorithms and approaches to doing quantum chemistry with deep learning according to scientific principles. Abbreviations in the table head used: Ψ - how the wave function is evaluated, OA — open access, PB — published before preprint release of this work, BOA — method needs the Born-Oppenheimer approximation, CA — method reaches chemical accuracy on most candidates, QC — method has edge over quantum-chemical methods. Abbreviations in the system size column: atm — atoms, sm — small molecules, smc — supramolecular complexes**.

| Method | Method | $\Psi$ | Chip | OA | Code | Commercial supp. | PB | BOA | System Size | CA | QC |
| --- | --- | --- | --- | --- | --- | --- | --- | --- | --- | --- | --- |
| Pauli Net | DNN | expl. | multi GPU | no | no | (Microsoft AI4Science) | yes | yes | atm, sm | no | no |
| DeepErwin | DNN | expl. | GPU | yes | no | no | yes | yes | atm, sm | no | no |
| DTNN | DNN | expl. | CPU | yes | yes | no | yes | yes | sm | no | no |
| SchNorb | GNN | expl. | GPU | yes | yes | no | yes | yes | sm | no | no |
| FermiNet | GNN | expl. | GPU | yes | yes | Google DeepMind | no | yes | atm, sm | no | no |
| HiMol | GNN | impl. | GPU | yes | yes | no | no | no | sm, smc | no | no |
| PDF | GNN | impl. | — | yes | no | Huawei | no | no | sm, smc | no | no |
| CARATE | GNN | impl. | CPU | yes | yes | no | — | no | sm, smc | yes | yes |

**Table 4.** Classification performance of CARATE as compared to other state of the art GNNs and kernel classifiers. Reference results were taken from *Morris et al*. ^44^.

| Method | MCF-7 | MOLT-4 | YEAST | Enzymes | Proteins |
| --- | --- | --- | --- | --- | --- |
| CARATE | 97 ± 2 | 95.4 ± 0.9 | 89.4 ± 0.4 | 95 ± 9 | 92 ± 9 |
| 1-WL | 94.5 | 94.6 | 89.2 | 50.8 | 72.6 |
| WL-OA | — | — | 88.2 | 56.4 | 73.4 |
| GR | 91.7 | 92.1 | 88.2 | 29.5 | 71.6 |
| SP | 91.7 | 92.1 | 88.3 | 39.3 | 75.6 |
| GIN-e | 92 | 92.4 | 88.3 | 38.7 | 72.2 |
| GIN-e-JK | 91.8 | 92.2 | 88.2 | 39.3 | 72.2 |

The Table 4 shows methods that aim to learn wave functions explicitly, like PauliNet ^38^, DeepErwin ^53^, DTNN ^33^, SchNorb ^52^, and FermiNet ^54^ as well as methods that aim to learn wave functions implicitly, like HiMol ^20^, PDF ^55^, and the algorithm used in this work, CARATE.

Around half of the methods were in work or completed before publishing the preprint of this present report. The other half was developed after the publishing of the first preprint. Work that was released after the first preprint of this work includes the explicit method FermiNet, as well as the implicit methods HiMol and PDF.

The two implicit methods model different aspects of the encoder structure in different ways. To investigate whether CARATE is a physically sound method, which can maintain state-of-the-art performance even two years after its first release, CARATE was also compared to the newer algorithms FermiNet, PDF, and HiMol.

HiMol is trying to mimic the concept of graph convolutions through the distance-based clustering algorithm ^20^. The PDF method aims to facilitate a generalization over spectral graph convolutions using a Fourier transform to tackle the problem of wave-function learning. Yet the method applies the same foundational mathematical concepts as the foundational GCN by just parametrizing the singular value decomposition of the Laplacian ^55^.

Most methods except for DTNN and CARATE recommend using GPUs, whereas HiMol can be trained in a reasonable amount of time on a CPU as well. Most methods have no commercial interests behind their research. However, PauliNet, FermiNet and PDF have ties to big tech companies either by being directly developed at those companies or in receipt of funding from a tech company at the time of writing the manuscript.

All explicit methods use the BOA as well. None of the implicit methods rely on the BOA. The system size is very limited for explicit methods stopping at the small-molecule scale. All implicit models can go up to supramolecular complexes.

All methods, except for CARATE, fail to achieve chemical accuracy across the datasets studied. Therefore, the conclusion is that only CARATE offers an edge in terms of speed and accuracy to classical quantum chemical methods.

Comparing CARATE to other state-of-the-art models in the realm of representation learning in quantum chemistry reveals that CARATE is the only algorithm that has an edge over quantum chemical simulations.

Moreover, the multi-headed self-attention mechanism allows efficient learning and prediction in low-data regimes. Besides, the CARATE algorithm may detect flawed data in given datasets or uncover statistically unrelated data, thus giving hints on where to improve the quality of the data in a data lake.

In contrast, in the highly distorted data regime, the CARATE algorithm may be inferior to classical methods based on supported-vector machines or random-forest classifiers. The behaviour, however, is expected, as GAT units are inserted flexibly to detect chemical patterns.

If the important pattern is not well represented in the dataset, the attention mechanism may forget the pattern due to the dropout probability of 0.6. To overcome this issue, highly distorted datasets can be modelled by adjusting the learning or dropout rate and training for more epochs to achieve similar to or better than baseline results.

In fact, the performance of the algorithm might also be increased by modifying the architecture in subsequent work via adjusting the network stack, using batch normalization or a slightly different dropout rate or dropout localization. However, the ablation study strongly indicates that the current network structure is crucial to the exceptional performance of the algorithm.

The ALCHEMY dataset poses a multitasking problem, and ZINC only predicts a single-class problem. Thus, the method generalizes to many operator problems from equation 7.

The MAE is used in the general Hartree-Fock scheme, making it a physically sound metric. Yet convergence is also achieved using the MSE error metric. Indeed, the convergence behaviour appeared to be more stable during the experiments.

## 7 Conclusions

The accurate results on quantum chemical datasets indicate that the algorithm is learning a wave function *ψ* to solve the Schrödinger equation via the last regression layer, such that the general Eigenvalue equation

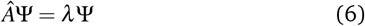

can be applied to understand the calculation. When considered with multitask problems, the network solves

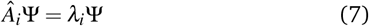

The experiments indicate that *A*_i_ can be a non-linear operator, as the activation function is the Leaky RELU function for the GAT units ^22^.

Thus, the network solves the generalized form of equation 2. The high accuracy for most of the classification and regression tasks suggests that the multi-head GAT units are decent encoders for molecular structures.

When considering the initial motivation of solving the eigen-value problem of an operator acting on a wave function, we must conclude that the encoding of the molecule through the CARATE unit indeed represents a wave function and the operator acting on the wave function is represented by the final classification or regression layer of the network.

The principle must by definition be self-replicating such that one can add layer after layer and this would still satisfy the eigen-value equations. Especially when presented with a descriptive data regime, CARATE shows low training times while achieving high accuracies for biomolecules and small molecules alike.

The idea of CARATE is similar to the one behind SchNet ^33^ but incorporating a self-interaction term and global attention such that high accuracies can be achieved. By incorporating more physical knowledge into the model, the encoding was framed as a Markov graph in the wider sense.

Even though there is a high dropout probability and one dropout layer is stacked as the last layer, accuracies are high.

On the other hand, it is known that large neural networks can model implicit attention ^56^, and thus the explicit attention mechanism models the investigated problems sufficiently. By modelling the multi-headed self-attention explicitly, the algorithm becomes intentionally more accurate and efficient.

Simulations utilizing CARATE seem promising. However, to verify the results, in later studies more experiments are necessary. It would be interesting to see how CARATE performs classification on larger datasets of biomolecules and small molecules because the fitting abilities of the proposed algorithm appear to be very high on large datasets such as ZINC and ALCHEMY.

In a recent master’s thesis, the construction of extremely detailed wave functions via random forests to solve a multi-scale problem of biodegradability was investigated successfully, leading to high calculation accuracies ^57^. Thus, simulations or quantum-chemical calculations on large macroscopic scales using the principles discovered in this and subsequent works are feasible.

The GAT layer uses masked attention ^22^ such that at least two training epochs are needed for the numerical fit. The masked-attention mechanism is actually similar to the convolutional mechanism, such that only the direct neighbors are relevant for the attention mechanism.

The attention mechanism is a message-passing mechanism and the message needs to propagate at least once. On small molecules the fit is thus fast and fitted after mostly two epochs.

For larger graphs such as the ENZYMES and PROTEINS dataset, the message needs to travel for longer distance and thus the training takes more iterations.

This interesting fitting behaviour gives a first intuition into the similarity of dynamic simulations and the training of deep-learning algorithms on molecular graphs.

The fitting behaviour also suggests that the algorithm is coarsegraining by default and coarse graining dynamically, as CARATE can fit data sets of both large graphs and small molecular graphs with high accuracies.

Thus, CARATE is behaving in a scale-invariant way and shows fractal behavior, as the algorithm can act on any graph across multiple scales simultaneously, incorporating self-similarity by default. The algorithm may thus have mathematical connections to Rado graphs or fractal graphs in general.

The CARATE graph-based algorithm does not rely on the BOA or other approximations and can fit experimental data to high accuracies, too. Thus, the method of using wave functions that are neural-network encodings of molecular graphs are superior to classical *ab initio* methods, as the BOA reaches its explanatory limit already at diatomic transition-metal fluorides ^58^.

The wave function is not encoded in Euclidean space and does not rely on atomic coordinates. This is an interesting approach, but it gets rid of the BOA by design.

It remains an open question whether the experimental data from the small compound datasets (MCF-7, YEAST, MOLT-4) are noisy and therefore whether CARATE correctly misclassifies some of the samples it is given. However, experiments with wave functions obtained from random forests suggest that it is possible to identify mislabelled data points in experimental datasets ^57^.

In the end, models should be derived from experimental values rather than theoretical considerations, such that new theories and models derived from CARATE or similar algorithms trained on experimental data of high quality may give new insights into the theoretical foundations of chemistry.

To summarize, the key qualities of the developed software and algorithms are:

- The algorithm fits up to numerical accuracy on the ZINC and ALCHEMY tasks.
- The algorithm shows best-of-class fits even after two years for classification problems PROTEINS and ENZYMES up to 100% accuracy on certain splits.
- The software can run on edge devices.
- The algorithm has low training times.
- The algorithm is explainable on the basic theoretical level. Further, in terms of diving deeper into the theoretical foundations, there may be a risk that too much explainability can hamper the creativity of a chemist.
- The software does not require expert knowledge.
- The software produces reproducible algorithms.

## 8 Outlook

Based on the accurate regression results, the GNC-GAT encoding enables the subsequently stacked layers to model a given, well-described problem accurately.

Considering the success of the general theoretical mindset presented in this work, a more in-depth study of the theoretical implications of the mathematical relations described here is mandatory for deriving a more solid theoretical framework.

The study is being prepared under the working title *Understanding chemistry through explaining quantum-chemical calculations from graph neural-networks*.

This upcoming work aims to use methods from the community of explainable AI, to understand the theoretical foundations of the underpinning fitting behavior of CARATE.

Future work will also focus on applying CARATE to multi-scale simulation problems, many-class multitask learning, transfer learning ^22^, and flawed or fraudulent data detection in chemical datasets.

It is also a promising venue to investigate how CARATE and derived algorithms perform on excited-state dynamics, as this paper shows that CARATE does not rely on the BOA as a mandatory approximation. Therefore, CARATE could bypass the current limitations of excited-state dynamics introduced by methods of state-of-the-art quantum-chemical methods. It would also be interesting how CARATE performs compared to multi-reference methods.

## Conflicts of interest

There are no conflicts to declare. Although for researching the patent system, a patent was filed in 2021 in the US. The patent was removed in 2023 by their government and would have required refiling. The author refused to refile the patent, as he did not believe in the principle of patents anyway.

The work was entirely funded with private funds of the author that were earned through working in food delivery jobs.

It is also worth noting that some censorship happened, and some actors tried to suppress the research presented in this article in different ways.

The author was cut off from academic resources, raw data vanished from different infrastructures, the author was threatened and denied funding as well as publishing opportunities in the first two years after the initial round of experiments.

## Acknowledgements

Special thanks go to my family and friends, the Berlin AI/ML community — especially to all members of BLISS e.V. — and my mentors at various universities, companies, and research institutes in Berlin.

Without you, I would not be where I am today. Furthermore, the paper would not have advanced to this stage. The work started as a small project. Still, the potential to improve the current way of doing quantum chemistry presented itself at an early stage during the first experiments.

The more time passed, the more work was done to finally prove more related theorems through different approaches. Thus, I want to thank especially Prof. Wolber, Katrin Denziger, Dr. Amsel, and Prof. Kümmerer for their support of the related master’s projects. During the master projects, I had more time to study other perspectives on the relationship between deep-learning and quantum physics.

This work needed a lot of reflection and questioning, even though the results were clear right from the beginning.

During this time, the software advanced to a reproducible state and the paper is more complete than the first preprint I wrote back in 2021.

Finally, I want to thank Frank Elminowski for the copy-editing the work.

The context for making sense of the results happened through many projects and was frozen by publishing as much of my research as possible, even though it is mostly in preprint stage or using formats other than traditional research papers. Interested readers can refer to my Research Gate profile to obtain the documents.

A distortion might happen easily and especially during mid-2022 through to 2023, a lot of research in my wider academic circle was influenced and enriched through dialogues and discussions.

The whole theoretical and scientific implications of solving the eigenvalue problem related to wave functions via DL were not understood, neither was the relation to wave mechanics clear to the community at the time when the first experiments of this work were done.

There are many reasons for this, and one of them might be the ongoing censorship of certain works related to chemistry and graph neural networks. Plenty of publications were only available on preprint servers, and one may assume that these articles ran into the same problems.

